# Deep-learning-based fMRI decoding of real-world size for hand-held objects

**DOI:** 10.64898/2026.02.01.703176

**Authors:** Cui Lang, Kosuke Miyoshi, Masaki Takeda

**Author notes:** Correspondence: Masaki Takeda **Email:**. **Competing Interest Statement:** The authors declare no competing interests.

## Abstract

Real-world object size is a fundamental dimension of visual cognition, supporting effective interaction with the environment and object manipulation. However, neural mechanisms encoding size have largely been inferred from extreme size comparisons, leaving the neural representation of subtle size differences within a manipulable, "hand-scale" range poorly understood. Here, we applied a three-dimensional deep neural network (3D DNN) to decode real-world size from whole-brain fMRI data (N = 50) using objects that all fall within a graspable range. The 3D DNN successfully decoded subtle size differences, achieving predictive accuracies comparable to multivariate pattern analysis. Crucially, Guided Gradient-weighted Class Activation Mapping (Guided Grad-CAM) revealed that discriminative size information was not confined to the ventral occipito-temporal cortex but extended to a distributed network. Notably, these regions spatially overlap with the specific brain areas implicated in size-perception distortions following brain damage. Our findings suggest that the brain represents subtle variations in object size through a non-linear, distributed network that transcends the traditional visual hierarchy. Specifically, this system likely orchestrates the integration of visual properties with semantic scaling and the multimodal convergence of vision, space, and memory.

## Introduction

Humans can often rapidly and automatically infer an object’s real-world size from visual input (Konkle & Oliva, 2012a), even though the same object’s retinal projection can vary substantially with viewing distance, viewpoint, and contextual background (Maltz et al., 2021; Sperandio & Chouinard, 2015). This ability is crucial for interacting with the environment, guiding action decisions such as grasping, carrying, and avoidance (Ansuini et al., 2015; Klein et al., 2020; Sburlea et al., 2021; Seegelke & Wühr, 2018). Beyond motor interaction, real-world size supports scene understanding (Meese et al., 2023), and serves as a foundational dimension for organizing and retrieving higher-level semantic representations (Hagen et al., 2024; Lu & Golomb, 2025).

A substantial body of research has demonstrated that information related to real-world size exhibits a reproducible spatial organization along the ventral visual pathway, particularly within the occipito-temporal cortex (OTC). Neural responses to large objects are typically biased toward relatively medial OTC regions, whereas neural responses to small objects are more prominent in relatively lateral, object-selective regions. At a macroscopic scale, this dissociation manifests as a continuous gradient along the medial–lateral axis of the OTC (Grill-Spector & Weiner, 2014; Huang et al., 2022; Julian et al., 2016; Konkle & Caramazza, 2013; Konkle & Oliva, 2012b). Together, these findings suggest that real-world size is not merely a semantic label assigned behaviorally, but is systematically encoded in the higher-level visual cortex, constituting an important dimension that structures object representations.

However, existing research on the neural mechanisms of real-world size has predominantly relied on comparisons between objects that differ drastically in ecological scale (Konkle & Caramazza, 2013; Konkle & Oliva, 2012b). In such paradigms, large objects are typically massive, immovable, and tied to navigational demands or scene structure, whereas small objects are often graspable tools or household items. Consequently, these studies are often confounded by concurrent variations in scene structure, action demands, and functional properties. As a result, it remains unclear to what extent the previously observed size-related neural organization is driven by extreme differences in scale per se, or whether it generalizes to a more constrained stimulus space primarily composed of manipulable everyday objects. Within this framework, a critical complementary question is whether brain activity patterns can still reliably distinguish between small and large conditions when objects are restricted to a relatively narrow size range with more subtle size differences.

In recent years, deep learning-based decoding of functional magnetic resonance imaging (fMRI) data has provided a complementary analytical framework for examining discriminative information embedded in distributed patterns of brain activity (Dong et al., 2018; Frey et al., 2021; Horikawa & Kamitani, 2017; Watanabe et al., 2023). Unlike univariate analyses that primarily focus on mean effects, and MVPA approaches that often assume linear separability, deep learning models can in principle integrate more complex spatial patterns across widespread brain regions. They therefore hold promise for providing complementary evidence of distributed information that may not be fully captured by traditional analytical approaches. Furthermore, when combined with interpretability methods such as Guided Grad-CAM (Selvaraju et al., 2017), deep decoding allows us to go beyond simple classification to identify precisely which voxel patterns drive decoding performance, thereby linking predictive accuracy directly to spatial neural evidence.

In this study, we examined fMRI data from 50 participants to investigate whether brain activity patterns remain discriminable between small and large conditions when stimuli are restricted to a relatively constrained set of manipulable everyday objects with subtle differences in size (Figure 1). We adopted a complementary analytical framework utilizing univariate analysis and MVPA as benchmarks, while introducing a three-dimensional deep neural network (3D DNN) to assess whole-brain decodability, integrated with Guided Grad-CAM interpretability mapping (Selvaraju et al., 2017) to localize the discriminative voxel contributions underlying the decoding performance (Figure 2). Our core questions were as follows: (1) whether whole-brain fMRI patterns can still discriminate between small and large conditions when stimuli are restricted to a relatively constrained set of manipulable objects; and (2) whether deep learning approaches can serve as a complement to conventional analyses, providing additional insights into these distributed discriminative patterns.

**Figure 1.**
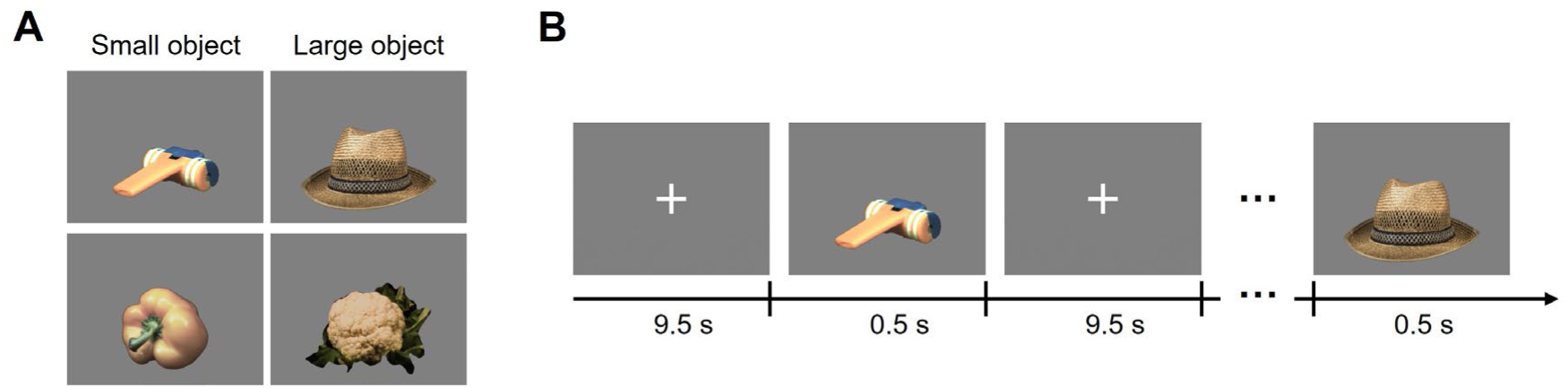
Visual stimuli and fMRI task paradigm. (A) Representative visual stimuli. (B) Experimental tasks and procedures. Participants performed a visual categorization task during fMRI scanning. Each run began with a 30-s fixation period, followed by 50 trials, including 40 stimulus trials and 10 blank trials. Each trial consisted of a 500-ms stimulus presentation and a 9,500-ms fixation interval. In the subsequent analyses, only correct object trials were included and were further classified as small-object or large-object trials according to their real-world size.

**Figure 2.**
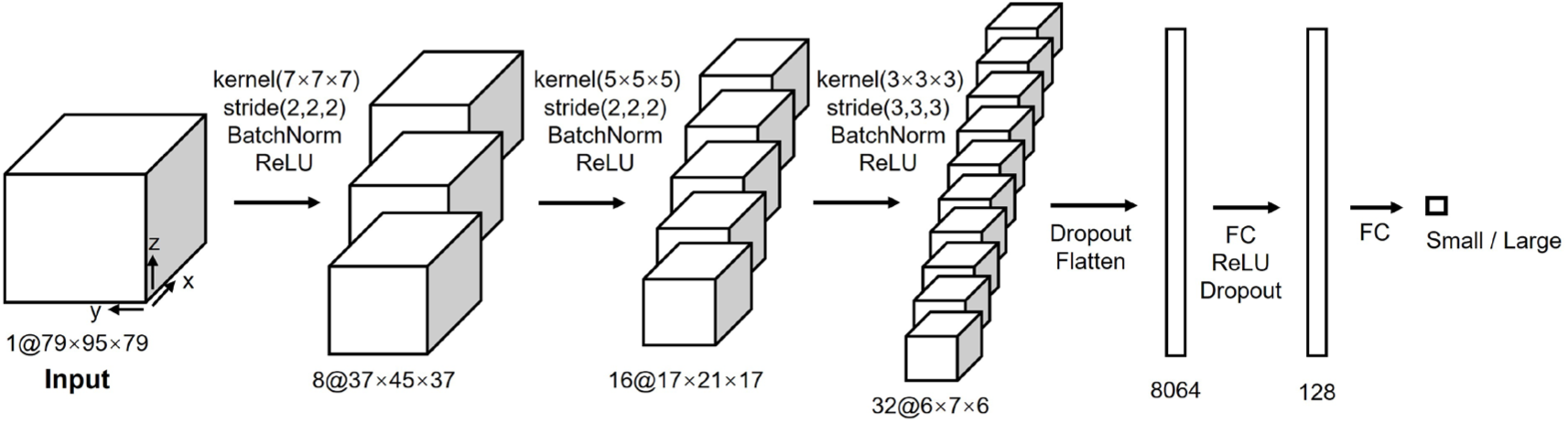
DNN model for fMRI data. The fMRI classifier was implemented as a 3D convolutional neural network consisting of three stacked 3D convolutional layers, followed by two fully connected layers that produced the final binary classification (small vs. large).

## Results

### Behavioral results

Fifty participants demonstrated high overall performance in the visual categorization task, with accuracy approaching ceiling levels (Figure 3). Crucially, behavioral performance was comparable between the two real-world size conditions. No significant differences were observed between the small-object and large-object conditions in terms of either accuracy (Wilcoxon signed-rank test, *P* = 0.255) or reaction time (*P* = 0.432).

**Figure 3.**
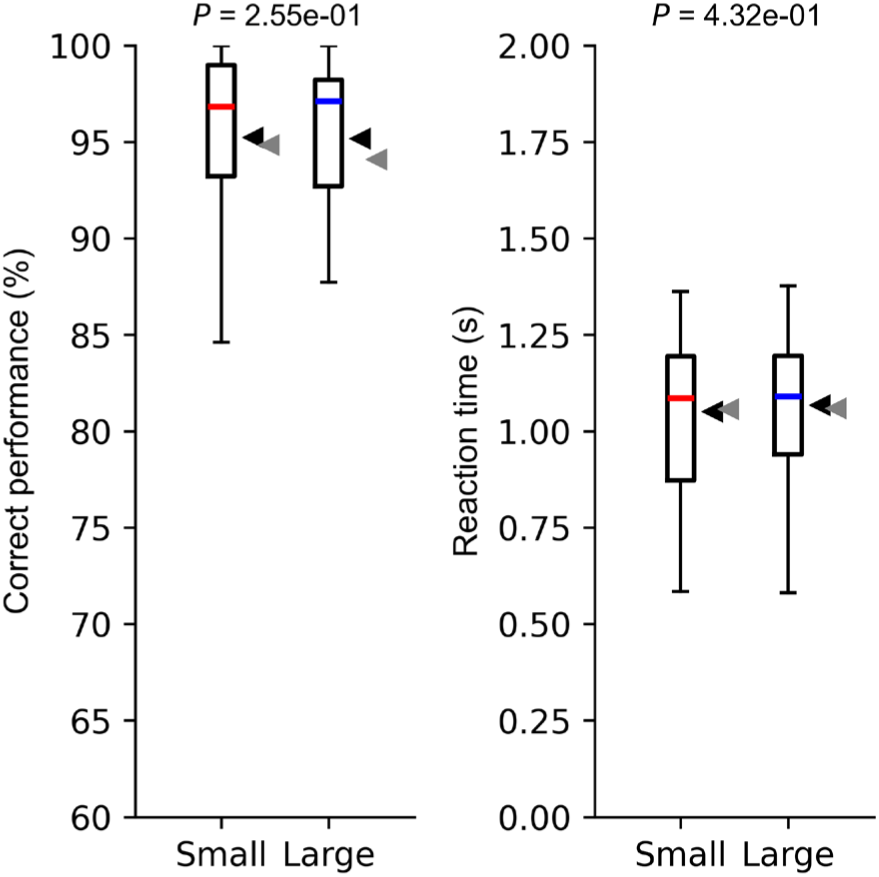
Behavioral performance for small and large object conditions. Comparisons of correct performance (left) and reaction time (right) are shown. No significant differences were observed between small and large object conditions for either metric (Wilcoxon signed-rank test; correct performance, *P* = 0.255; reaction time, *P* = 0.432). To illustrate temporal stability, black and gray arrowheads indicate mean values for the first and second halves of the trials, respectively.

To examine potential practice or fatigue effects over the course of the experiment, we divided each participant’s trials into the first and second halves chronologically. Neither behavioral performance nor reaction times differed significantly between the first and second halves across conditions (Wilcoxon signed-rank test, behavioral performance: *P* > 0.57; reaction time: *P* > 0.20), indicating that behavioral performance remained stable throughout the experiment.

Taken together, these results confirmed that the small-object and large-object conditions were well matched in terms of task difficulty. Furthermore, practice- or fatigue-related confounds appear minimal and therefore unlikely to introduce systematic bias in the subsequent neuroimaging analyses.

### Spatial distribution of real-world size preference in univariate fMRI

In the univariate fMRI analysis, we compared BOLD responses between the small-object and large-object conditions to identify brain regions sensitive to real-world size. For the “small > large” contrast, no brain regions survived whole-brain FWE correction (*P* < 0.05), indicating a lack of robust activation favoring small objects. In contrast, the “large > small” comparison revealed multiple significant clusters, primarily located in bilateral posterior visual cortex and ventral occipitotemporal regions (Figure 4A). These regions encompassed extensive portions of the occipital lobe and the occipitotemporal junction, indicating that large-object stimuli elicited significantly stronger BOLD responses in these visual regions compared to small objects. Crucially, as low-level visual features were matched between conditions, these differential activations likely reflect higher-level size processing rather than basic visual confounds. Meanwhile, the analyses of the “small-object vs. baseline” and “large-object vs. baseline” contrasts further showed that both conditions, relative to baseline, elicited significant responses across extensive visual cortical regions as well as parts of higher-order associative cortex, with broadly similar spatial distributions overall (Figure S2). These results indicate that both the small-object and large-object conditions reliably engaged the visual system. Overall, these univariate results suggest that at the coarse scale of mean regional activation, real-world size information is predominantly manifested as a large-object preference, whereas voxel-wise preference for small objects is comparatively weak.

**Figure 4.**
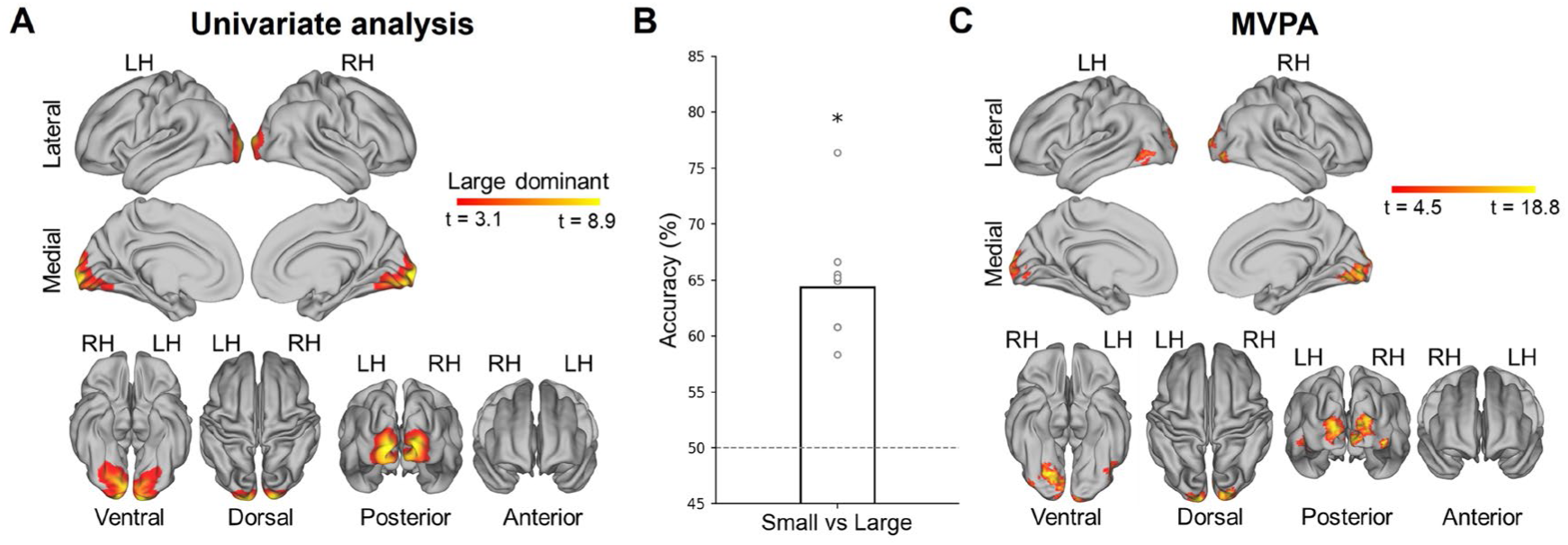
Spatial representation of real-world object size revealed by univariate analysis and MVPA. (A) Univariate activation maps. Hot color denotes significant clusters exhibiting stronger responses to large objects (“large > small”). Note that no significant clusters were detected for the contrast of “small > large”. (B) Whole-brain MVPA decoding performance. The bar denotes mean classification accuracy; dots represent accuracy for individual cross-validation models. The dashed line represents chance level (50%). An asterisk denotes significant deviation from chance (t-test, FDR-corrected *q* < 0.05). (C) Searchlight MVPA information map showing regions with significant local decoding accuracy.

### Spatial distribution of real-world size in MVPA

Whole-brain MVPA was performed using all brain voxels as features to classify small versus large object conditions via a linear SVM with 9-fold cross-validation. The classification across the nine folds yielded a mean accuracy of 64.39% (Figure 4B). This decoding performance was significantly above chance (t-test against chance level, *P* < 0.05), indicating that distributed fMRI patterns across the whole brain contain stable and decodable information regarding real-world size.

We next performed searchlight MVPA to spatially localize specific brain regions encoding real-world size information. Significant decoding clusters were identified across multiple regions, indicating that local voxel patterns reliably discriminated between the small and large object conditions. These clusters were primarily located in bilateral occipital cortex and ventral occipitotemporal regions (Figure 4C).

Overall, the searchlight results exhibited a spatial topography highly consistent with the univariate findings, reaffirming the critical role of bilateral posterior visual cortex and occipitotemporal regions in representing the real-world size of objects. Furthermore, these results demonstrate that size information is encoded in distributed local multivoxel patterns extending beyond the mean activation differences captured by univariate analysis.

### Deep-learning-based decoding of real-world size of objects using a 3D DNN

In the deep-learning analysis, we trained a three-dimensional deep neural network (3D DNN) to classify real-world object size using whole-brain fMRI voxel patterns. Figure 5A shows the decoding performance on the test set as a function of the number of averaged trials. At the single-trial level, mean accuracy was only marginally above chance (mean, 51.20%; range, 48.90–53.30%). However, averaging three or more trials led to a monotonic increase in decoding accuracy, and for all averaging conditions, accuracy was significantly above chance (t-test with FDR correction, *q* < 0.05). With nine-trial averaging, the mean decoding accuracy exceeded 65.75% (best: 69.51%), which was comparable to that achieved by whole-brain MVPA (mean: 64.39%). These results indicate that the deep-learning decoder can leverage real-world object size–related information, and that trial averaging effectively enhances the signal-to-noise ratio of the fMRI patterns.

**Figure 5.**
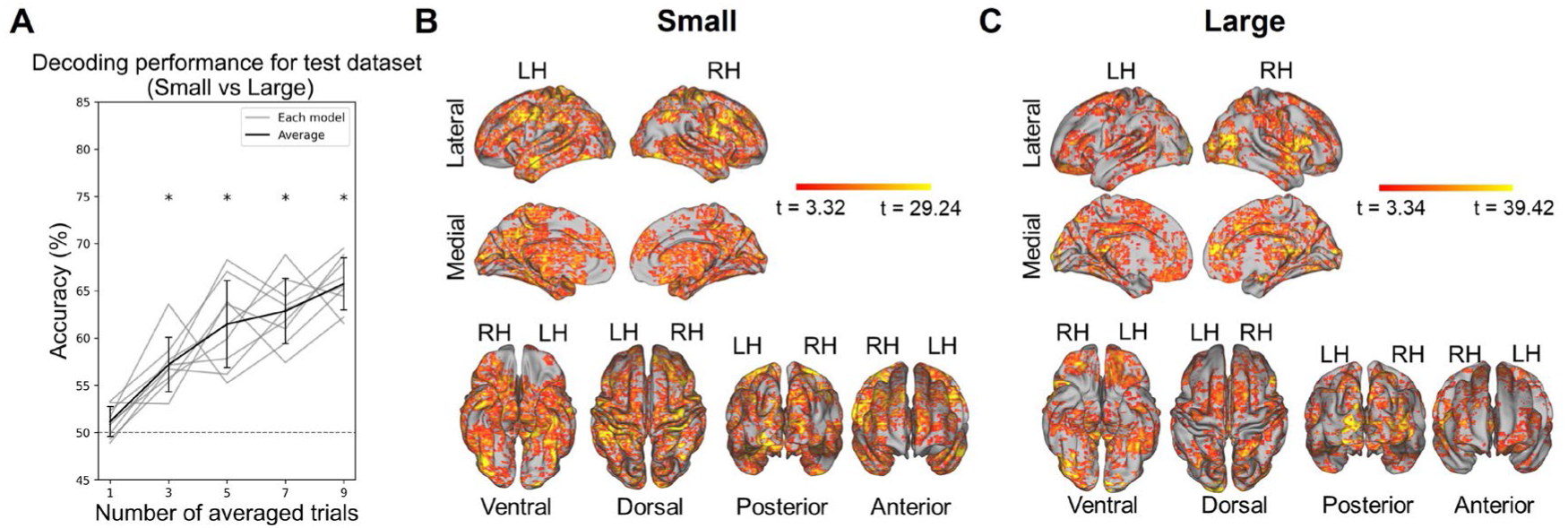
Spatial representation of real-world objects size revealed by deep learning. (A) Decoding performance on the independent test dataset as a function of the number of averaged trials. The thick line indicates the mean classification accuracy; thin lines represent the accuracy in each model. *, FDR corrected *q* < 0.05 (t-test for accuracy distribution against chance level). (B) Spatial organization of the Guided Grad-CAM values for small objects. The threshold of significance was set at *P* < 0.05 corrected with cluster-wise FWE based on a non-parametric permutation test. (C) Corresponding saliency maps for large objects.

To identify the brain regions driving the model’s predictions, we computed class activation maps using Guided Grad-CAM for the best-performing model and projected them onto the standard cortical surface (Figure 5B–C). Discriminative features were primarily distributed across bilateral occipital cortex and ventral occipitotemporal regions, extending into parts of parietal and frontal cortex. Notably, the size-related saliency patterns for small and large objects were spatially widespread and interleaved across the cortical surface, rather than being confined to a few isolated nodes.

To quantify these regional differences, median Guided Grad-CAM values were summarized within five anatomical divisions (Figure 6). Compared with the large-object condition, the small-object condition exhibited higher attribution values across all divisions, with the most pronounced differences observed in the parietal and occipital divisions. This pattern suggests that, within the classification model used in the present study, regions associated with the occipital and parietal cortices contribute more strongly to the spatial evidence supporting small-category decisions. Correspondingly, attribution values in these divisions were relatively lower in the large-object condition, indicating that the model’s discrimination of the large category does not rely on these regional spatial patterns to the same extent under the current task.

**Figure 6.**
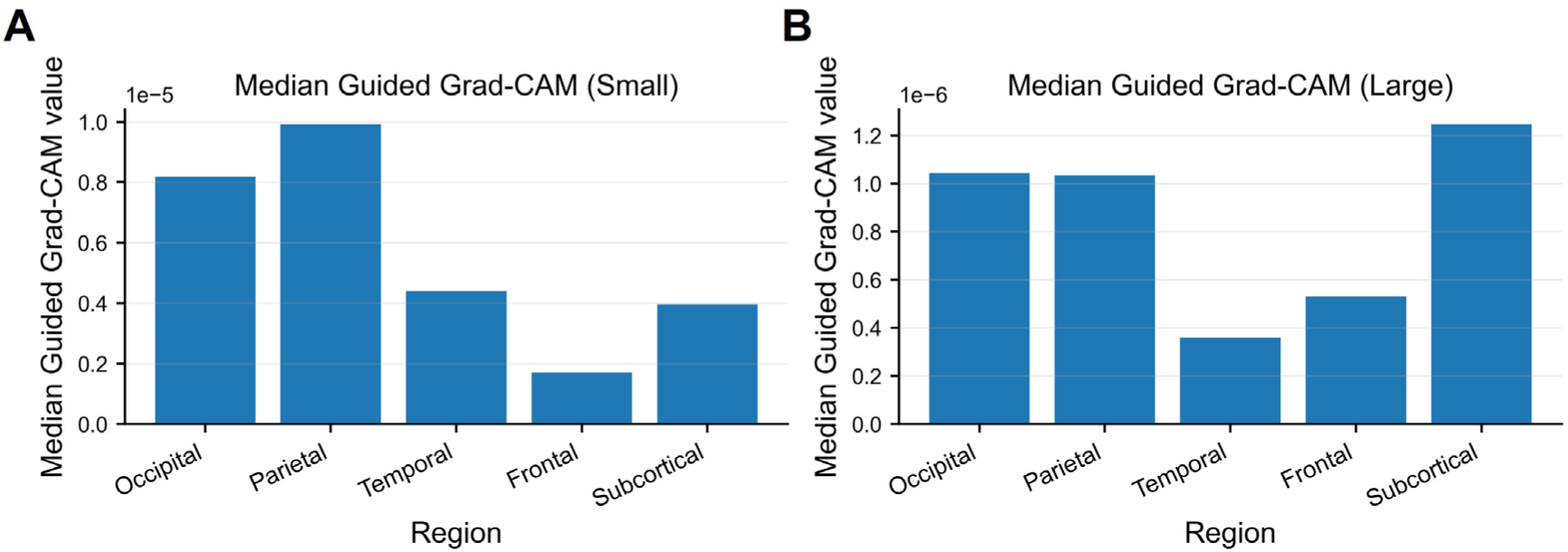
Regional distribution of Guided Grad-CAM values. (A) Median Guided Grad-CAM values summarized within five major anatomical divisions for the small-object condition. (B) Corresponding values for the large-object condition. Note that the vertical axes for the two panels are scaled differently.

Among 110 cortical and subcortical ROIs, 102 ROIs were identified as significant for the small-object condition and 85 for the large-object condition (Tables S1–S2), indicating that these ROIs tended to contribute positively to the target class score. To further illustrate the key regions within each anatomical division that contributed most strongly to the model’s discrimination, we selected the top five ROIs ranked by Guided Grad-CAM values within each anatomical region (Figure 7). In the small-object condition, dominant contributions in the occipital region were concentrated in the lateral occipital cortex and early visual cortex–related regions (Figure 7A). High-contributing ROIs in the parietal region were primarily distributed in the superior parietal lobule, and parietal operculum, as well as the precuneus and angular gyrus, the latter two of which are situated within the temporo-parietal/occipito-parietal junction. In the temporal region, high-contributing ROIs were mainly located in the posterior part: the posterior inferior temporal gyrus, posterior fusiform gyrus, and posterior parahippocampal gyrus. In the frontal region, high-contributing ROIs included the frontal pole, superior frontal gyrus and juxtapositional lobule cortex (the supplementary motor cortex). Subcortically, the top-ranked ROIs predominantly involved the insula, striatum (putamen/caudate), pallidum, and hippocampus.

**Figure 7.**
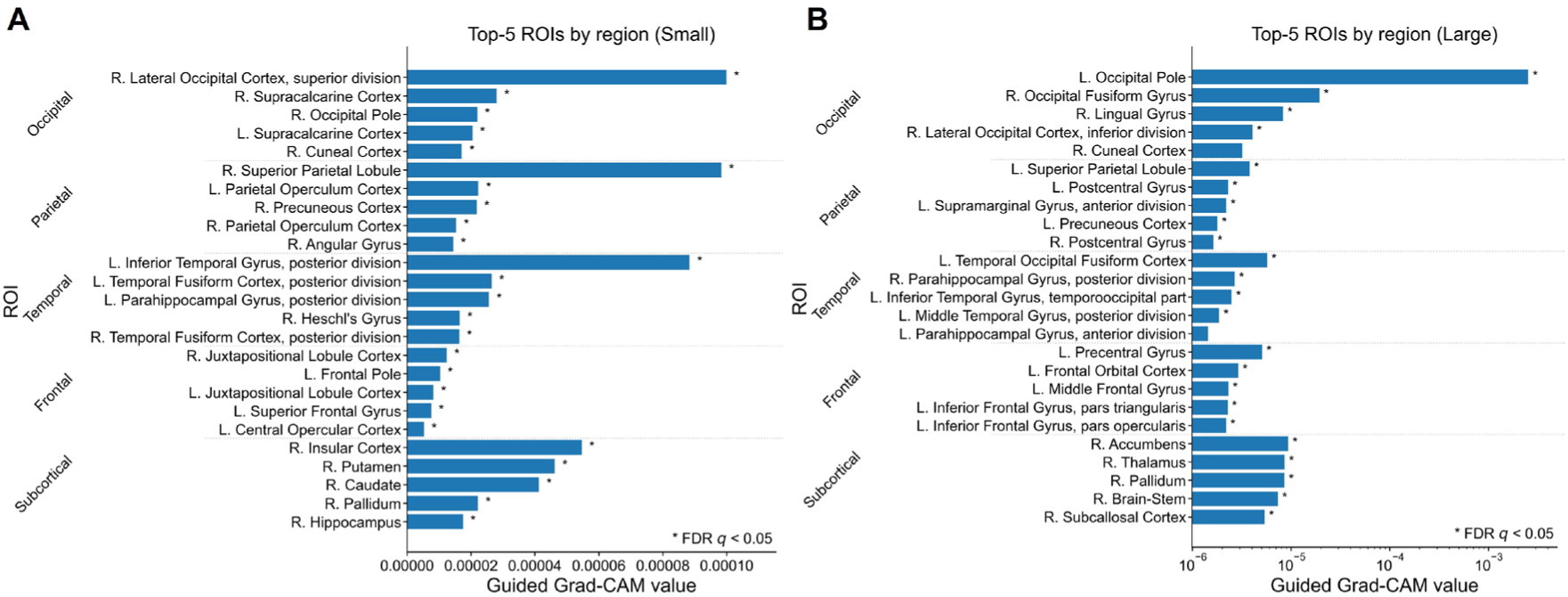
Top-ranked ROIs contributing to size decoding. (A) The five ROIs with the highest mean Guided Grad-CAM values within each anatomical region for the small-object condition. Asterisks denote ROIs showing significant effects after FDR correction (*q* < 0.05). (B) Corresponding top-ranked ROIs for the large-object condition. Note that the x-axis is displayed on a logarithmic scale to accommodate the wide dynamic range of Guided Grad-CAM values. Asterisks denote ROIs showing significant effects after FDR correction (*q* < 0.05).

By contrast, in the large-object condition (Figure 7B), the occipital region still contained the most prominent top-ranked ROIs (e.g., occipital pole, occipital fusiform gyrus, and lingual gyrus). In the parietal region, top-ranked ROIs were found in the superior parietal lobule and postcentral gyrus, as well as the supramarginal gyrus and precuneus, the latter two of which are situated within the occipito-parietal junction. Temporal contributions were primarily centered on the fusiform and parahippocampal/inferior temporal subdivisions near the temporo-occipital junction. In addition, multiple significant ROIs emerged in the frontal region (precentral gyrus, orbitofrontal cortex/middle/inferior frontal gyrus). Subcortically, top-ranked ROIs included the nucleus accumbens, thalamus, pallidum, brainstem, and subcallosal cortex.

Taken together, these results suggest that real-world size information is not only represented along the occipito-temporal visual pathway but may also be further integrated within a distributed network spanning parietal–frontal–subcortical systems, thereby supporting higher-order cognitive processing.

## Discussion

This study establishes that the brain encodes real-world size information for manipulable objects through a distributed network extending far beyond the canonical visual cortex. By combining deep-learning-based decoding with interpretable mapping, we demonstrate that even subtle size differences within a "hand-scale" range are robustly represented in whole-brain fMRI patterns.

### Distinct and shared neural representations of real-world size for small and large objects

While univariate analysis confirmed a macroscopic bias for large objects in ventral temporal regions, it failed to capture the fine-grained representations of small objects (Figure 4A). Our deep learning results not only overcame this limitation but also revealed multivariate patterns encoding size information that are distributed across the whole brain (Figure 5B-C), which could not be captured even by MVPA (Figure 4C).

Guided Grad-CAM visualization uncovered a size-dependent asymmetry in how the brain represents size. Recognition of relatively small objects within a hand-scale range preferentially recruited the occipito-parietal pathway [e.g., superior parietal lobule (SPL), supramarginal gyrus] (Figure 6A). Conversely, relatively large objects within a hand-scale range relied more heavily on ventral stream regions and subcortical structures (Figure 6B). While all objects were graspable, small objects (e.g., whistle) typically require fine motor control (precision grip) (Hussain et al., 2024; Van Polanen et al., 2022), whereas large objects (e.g., hat) involve bimanual or coarser handling. This difference in type of manipulation likely drove the parietal activity in the small-object condition.

Notably, significant brain regions identified for both conditions were concentrated within occipitotemporal visual regions, closely aligning with the size-sensitive visual regions identified by searchlight MVPA. Furthermore, the extensive involvement of parietal areas (e.g., superior parietal lobule, precuneus, parietal operculum, and angular gyrus), frontal regions (e.g., frontal pole, superior and middle frontal gyri, and paracentral lobule cortex), and subcortical structures (e.g., insula, striatum, globus pallidus, thalamus, and hippocampus) suggests that real-world size processing is embedded in coordinated processing across multiple systems, including attention, decision-making, action preparation, and memory/value-related computations (Branzi et al., 2022; Fairchild et al., 2025). It is noteworthy that real-world size information was robustly decoded even though the participants performed a visual categorization task rather than an explicit size-judgment task. This finding strongly aligns with previous behavioral and neuroimaging evidence demonstrating that real-world size is an automatic and obligatory property of object representation (Hagen et al., 2024; Konkle & Oliva, 2012a). Once an object is visually recognized, its real-world scale is spontaneously accessed to support potential motor interactions and spatial scaling, thereby allowing the deep-learning decoder to capture this task-independent size representation.

### The "common network" hypothesis: linking healthy perception to AIWS

Perhaps the most significant finding of this study is the systematic overlap between the discriminative regions identified by our model and the lesion network map of Alice in Wonderland Syndrome (AIWS), a neurological condition characterized by distortions in visual perception (metamorphopsia) and the self-body schema (Friedrich et al., 2024; Piervincenzi et al., 2022, 2023). Prior studies have frequently suggested that damage to the temporo-parietal-occipital (TPO) junction—including the extrastriate body area and inferior parietal cortex—as well as the SPL, causes AIWS.

As shown by the top-ranked ROIs for each anatomical region in Figure 7, our ROI analysis successfully identified the distinctive contribution of the TPO junction across its parietal (angular/supramarginal gyrus), temporal (posterior middle temporal gyrus and temporooccipital inferior temporal gyrus), and occipital (lateral occipital cortex) components, as well as the SPL. These brain regions are known to play a crucial role in semantic scale judgment and the multimodal integration of vision, space, and memory, and classic AIWS symptoms such as micropsia and macropsia are often discussed in terms of abnormal calibration of “scale mapping” or reference-frame alignment at such integrative nodes (Friedrich et al., 2024; Piervincenzi et al., 2022). Thus, the current method, which enables the highly sensitive visualization of individual voxel contributions, has the potential to facilitate the early detection of neurological conditions such as AIWS.

### Limitations and Future Directions

First, the present study is based on an existing dataset in which the stimuli were limited to four categories of manipulable objects. Future work is therefore needed to examine the extent to which the present findings generalize to more comprehensive representations of real-world size using datasets with a broader range of object categories.

Second, we acknowledge that the 3D DNN required trial-averaging to overcome the low SNR of single-trial data and achieve predictive accuracies comparable to linear MVPA. Thus, the primary advantage of our 3D DNN approach lies not in maximizing raw classification accuracy, but rather in its capacity to capture non-linear, spatially distributed feature representations.

Future studies may further evaluate the robustness and generalizability of the present findings by employing more strictly controlled stimuli (e.g., controlling for low-level visual features as well as semantic and action-related factors), and by conducting cross-dataset validation.

## Materials and methods

### Dataset

The present analyses were performed on an existing fMRI dataset collected and reported by Watanabe et al. (2023). All MRI data were acquired using a Siemens 3T Prisma scanner. Functional images were obtained using a gradient-echo echo-planar imaging (EPI) sequence with the following parameters: repetition time (TR) = 2,000 ms, echo time (TE) = 27 ms, flip angle (FA) = 90°, 36 contiguous slices, slice thickness = 3 mm, and an in-plane resolution of 2 × 2 mm². Each functional run consisted of 265 volumes. The first five volumes were discarded from subsequent analyses to allow for equilibration of longitudinal magnetization.

High-resolution T1-weighted anatomical images were acquired using a magnetization-prepared rapid gradient echo (MP-RAGE) sequence with the following parameters: TR = 2,400 ms, TE = 2.32 ms, FA = 8°, 192 slices, slice thickness = 0.9 mm, and an in-plane resolution of 0.93 × 0.93 mm².

### Participants

The original cohort comprised 53 right-handed participants (34 males, 19 females; aged 18-25 years), all of whom reported normal or corrected-to-normal vision (Watanabe et al., 2023). Following the quality control and behavioral criteria established by Watanabe et al., three participants were excluded from the final analyses (two participants whose accuracy fell below 80% in at least one subcategory, and one participant whose mean reaction time exceeded 1.5 s). Consequently, data from 50 participants were included in the subsequent analyses. No new data were collected. This experiment was approved by the Institutional Review Board of Kochi University of Technology. All participants provided written informed consent prior to the experiment.

### Stimuli

In the original study, participants completed a visual categorization task. The stimulus set consisted of 40 color images: 20 face images and 20 object images. Face images were drawn from the ATR Facial Expression Database (DB99) and included two male and two female identities with neutral facial expressions; each identity was photographed from five different viewpoints. Object images were taken from the Amsterdam Library of Object Images (ALOI) (Geusebroek et al., 2005) and consisted of four object types: two natural objects (bell pepper, broccoli) and two artificial objects (hat, whistle), each captured from five viewpoints. During the experiment, participants viewed color images presented at the center of the screen (visual angle ≈ 7.4° × 5.8°), along with a central white fixation cross (0.3° × 0.3°). To reduce the confounding effects of low-level image statistics, image contrast and luminance were matched across all stimuli using the SHINE toolbox (customized via MATLAB) (Willenbockel et al., 2010).

For the present secondary analysis, we focused exclusively on object trials. Face trials were excluded, and the objects were regrouped into two conditions based on their typical real-world physical size—hereafter referred to as “small” and “large” (Figure 1A). Importantly, the typical real-world sizes of both small and large objects fall within a manipulable, peripersonal scale. They can be directly manipulated by the hands and clearly contrast with non-manipulable, navigable-scale objects. Thus, all stimuli in our analysis belong to the same broad functional category of graspable objects. We confirmed that low-level image statistics, including contrast, luminance, similarity to other stimuli of the same category, pixel intensity, and spatial frequency power distributions, were balanced across the size categories (Figure S1).

### Task procedure

In the original experiment, the task was programmed and administered using Psychtoolbox version 3 (http://psychtoolbox.org/). Participants performed a four-alternative forced-choice (4AFC) task, judging the visual subcategory of each stimulus (male face, female face, natural object, or artificial object) by pressing one of four buttons. At the beginning of each fMRI run, an instruction screen displayed the mapping between the subcategories and button positions. This mapping was pseudo-randomly varied across runs to decouple stimulus identity from motor response associations (Figure 1B).

Each run began with a 30-s fixation period, followed by the main task, which consisted of 50 trials. Of these, 40 trials presented a face or object image, and the remaining 10 trials presented a blank image to allow the blood-oxygen-level-dependent (BOLD) signal to return to baseline (Boynton et al., 1996; Jimura et al., 2016; Yamashita et al., 2009). Each trial lasted 10 s: a 500-ms stimulus presentation was followed by a 9,500-ms fixation interval.

Within each run, the 40 stimulus images were presented once in a pseudo-random order without performance feedback. Participants completed on average 9.96 ± 2.88 runs (496 runs in total). In the original study, trials with reaction times longer than 2 s (approximately 5% of all stimulus trials) were first excluded, and only trials with correct responses were entered into subsequent analyses.

As noted above, we focused exclusively on object trials. For each correct object trial, we re-labeled trials as a “small-object” or “large-object” trial according to the physical real-world size of the corresponding object. Ultimately, only correct object trials were included in the subsequent fMRI analyses.

### Behavioral data analysis

Behavioral data were analyzed using MATLAB (MathWorks, Natick, MA). For each trial, we extracted the stimulus type (small-object / large-object), response accuracy (correct or incorrect), and reaction time (RT), and computed behavioral indices for the small and large conditions at the individual-subject level. To reduce the influence of outlier responses, we excluded trials with RT > 2 s and retained only correct trials with RT ≤ 2 s for analysis. For the remaining correct trials, we calculated mean accuracy and mean RT for the small and large conditions for each participant. A Wilcoxon signed-rank test was used to compare these measures at the group level.

To examine potential practice or fatigue effects, we divided each participant’s trials into the first and second halves chronologically, and the behavioral indices were then aggregated into group-level summaries.

### fMRI preprocessing

fMRI preprocessing was performed following the pipeline reported by Watanabe et al. (Watanabe et al., 2023). The analysis was implemented using SPM12 (http://fil.ion.ac.uk/spm/). For each run, the initial five volumes were discarded to allow for longitudinal magnetization equilibration. Slice-timing correction was then applied to compensate for differences in acquisition time across slices, and head motion was corrected using rigid-body realignment, yielding six motion parameters (three translations and three rotations) for use as nuisance regressors in subsequent modeling. Functional images were then co-registered to each participant’s T1-weighted anatomical image and spatially normalized to the MNI152 standard space using a combination of linear and nonlinear transformations, followed by resampling to 2 × 2 × 2 mm³ isotropic voxels. A high-pass temporal filter was applied to the time series to remove low-frequency drifts.

To preserve fine-grained spatial structure relevant to deep-learning decoding, DNN analyses were performed on unsmoothed functional data, consistent with prior reports on decoding sensitivity to smoothing (Chen et al., 2016; Kamitani & Sawahata, 2010; Op de Beeck, 2010; Yang et al., 2025). Similarly, spatial smoothing was not employed in the MVPA analysis to avoid blurring domain-specific information and fine-scale patterns. In contrast, for the univariate general linear model (GLM) (Worsley & Friston, 1995) analyses, normalized functional volumes were smoothed with a 6-mm FWHM Gaussian kernel to enhance signal-to-noise ratio and satisfy the assumptions of voxel-wise statistical inference.

### Univariate analysis

We performed a standard univariate fMRI analysis (Matsui et al., 2022; Tanaka et al., 2020) using FSL (https://fsl.fmrib.ox.ac.uk/fsl/docs/) to test for BOLD amplitude differences between the small-object and large-object conditions. At the single-subject level, a GLM was fitted to the preprocessed volumes using the *FEAT* module. Correct trials for the small-object and large-object conditions were modeled as separate event-related regressors (duration = 0.5 s), convolved with a standard gamma-shaped hemodynamic response function (HRF). To account for potential confounds, nuisance regressors were included for six rigid-body motion parameters, incorrect trials, and physiological noise components (white matter, CSF, and respiration). Temporal autocorrelations were corrected using *FILM* prewhitening (Woolrich et al., 2001) and the voxel time series were high-pass filtered to attenuate slow drifts.

Contrast images comparing large-object versus small-object conditions were computed for each run and subsequently combined at the subject level using a fixed-effects model. Group-level statistical inference was performed using nonparametric permutation testing via FSL’s *randomise* tool (5,000 permutations) (Nichols & Holmes, 2002; Winkler et al., 2014). Statistical maps were thresholded using threshold-free cluster enhancement (TFCE) with a family-wise error (FWE) corrected significance level of *P* < 0.05 (Eklund et al., 2016). Finally, to facilitate comparison with MVPA and deep learning results, significant volumetric clusters were projected onto the midthickness cortical surface using Connectome Workbench.

### Multivoxel pattern analysis (MVPA)

To complement the univariate results with a multivariate perspective, MVPA was conducted to decode real-world object size from distributed neural patterns (Jimura & Poldrack, 2012).

#### MVPA implementation

Analyses were implemented in MATLAB (https://www.mathworks.com/) using The Decoding Toolbox (TDT; https://sites.google.com/site/tdtdecodingtoolbox/) (Hebart et al., 2015). A LIBSVM-based linear support vector machine (SVM) with a regularization parameter of C = 1.0 was employed as the classifier (Chang & Lin, 2011). Prior to classification, single-trial β images were normalized using global min‒max scaling, to minimize signal amplitude variability across trials. Consistent with the deep learning analysis, five participants were reserved *a priori* as an independent test cohort, while the remaining 45 constituted the pool for a 9-fold cross-validation scheme, where the small-object and large-object conditions were labeled as +1 and −1, respectively.

#### Whole-brain MVPA

Whole-brain binary classification was performed using TDT’s whole-brain decoding mode (*analysis = ’wholebrain’*). For each participant, single-trial β images were registered to MNI152 standard space, and all voxels within the standard MNI152 2-mm whole-brain mask served as features. Cross-validation followed a 9-fold leave-one-run-out scheme, with each fMRI run serving as one fold: in each fold, one run was used as the validation set and the remaining runs as the training set. Decoding accuracy, defined as the proportion of correctly classified trials, was averaged across the nine folds for each participant. At the group level, performance was assessed using a one-sample t-test against chance (50%).

#### Searchlight MVPA

To further localize brain regions carrying information about real-world object size, we conducted searchlight MVPA (Stelzer et al., 2013) using TDT in MATLAB. A spherical searchlight with a radius of 6 mm was defined around each voxel within the whole-brain mask. Local multivoxel patterns were extracted and used to train and test the linear SVM using the same 9-fold cross-validation scheme described above. The resulting searchlight accuracy maps, representing the local discriminability of the small-object versus large-object conditions, were subjected to group-level statistical inference. Significant clusters were identified using cluster-based permutation testing (cluster-forming threshold of *P* < 0.001, cluster-level threshold of FWE-corrected *P* < 0.05 ; 5,000 permutations) (Nichols & Holmes, 2002; Stelzer et al., 2013; Winkler et al., 2014).

### Deep learning

To further elucidate how the brain represents real-world object size, we implemented a deep learning-based decoder on the same fMRI dataset used in the univariate and MVPA analyses, performing binary classification between the small-object and large-object conditions. The deep neural network classifier was implemented in PyTorch, and the overall analysis pipeline followed the methodology described by Watanabe et al. (2023).

#### Classifier and data partitioning

We used single-trial fMRI data from all 50 participants. Five participants were reserved *a priori* as an independent test cohort, while the remaining 45 constituted the pool for model development. For hyperparameter tuning and validation we employed a subject-level 9-fold cross-validation within the pool: in each fold, a group of five participants served as the validation set and the other 40 as the training set. This partitioning ensures subject-wise independence across training, validation and final test evaluations.

For each hyperparameter configuration, we fitted one model per fold (nine models in total). Hyperparameter searches and model selection used only the training and validation splits; the held-out test participants were not accessed during model tuning. Preliminary testing identified three hyperparameters with strong influence on fMRI decoding: the trial-averaging factor (*N*), weight-decay, and the optimizer learning rate. We therefore ran a grid search where *N* ∈ {1, 3, 5, 7, 9}, the learning rate was sampled logarithmically between 1×10⁻⁴ and 1×10⁻² (five values), and weight decay was searched over five values between 0 and 1×10⁻².

After completing 9-fold cross-validation and identifying the optimal hyperparameter combination, we evaluated the final model on the independent test set. To remove class-imbalance effects on test performance, we randomly subsampled trials in the held-out test set so that the small-object and large-object classes were balanced (359 trials per class); this ensured a 50% chance level for binary classification.

#### Network architecture

The fMRI classifier was implemented as a 3D convolutional neural network consisting of three stacked 3D convolutional layers, followed by two fully connected layers that produced the final binary classification (“small-object vs. large-object”) (Figure 2) (Frey et al., 2021). Training was conducted using a batch size of 100 examples, the Adam optimizer (Kingma & Ba, 2017), and early stopping based on the validation loss. On top of this base configuration, the key hyperparameters—trial-averaging window size, learning rate, and weight decay—were optimized via the grid search described above.

Inputs to the network were single-trial 3D β-maps registered to standard MNI space and masked to exclude extracerebral voxels. Voxel intensities were normalized to preserve comparability across analyses.

#### Trial averaging

To increase signal-to-noise ratio (SNR), we applied an *N*-trial averaging procedure to correct object trials. For each participant and condition, trials were randomly permuted and repeated *N* times along the trial axis; the repeated sequence (length *T*×*N*, where *T* is the original trial count) was then partitioned into blocks of *N* trials. Voxel-wise means were computed within each block, yielding *T* averaged trials per condition.

This approach maintained the total number of samples equal to the original trial count, thereby facilitating direct performance comparisons across different averaging windows (*N* ∈ {1, 3, 5, 7, 9}). The averaging parameter *N* yielding the highest mean validation accuracy was selected for the final evaluation.

#### Visualization

In this study, we applied Guided Grad-CAM (Selvaraju et al., 2017) to visualize the input-space patterns on which the deep model relies during classification. This method integrates two sources of information: gradient-weighted class activation mapping and guided backpropagation. On the one hand, it uses the gradients of the target class output with respect to the final convolutional layer to obtain a coarse, class-specific response distribution. On the other hand, this class-relevant spatial response is combined with high-resolution gradient information at the input level, thereby producing saliency maps with enhanced spatial precision. For fMRI data, each voxel value in the resulting saliency map can be interpreted as the relative contribution of the local signal pattern at that location to supporting the model’s prediction for a given class. Larger values within a region indicate a greater tendency for that region to provide evidence favoring the corresponding class output. Unlike the standard Grad-CAM implementation which uses global average pooling (GAP), GAP was omitted in this study. This is because, in the context of three-dimensional convolutional networks, GAP substantially smooths gradient information, thereby weakening spatial localization and, in particular, compromising the preservation of fine-grained structural information in volumetric data. Accordingly, we adopted a strategy similar to that of Layer-CAM (Jiang et al., 2021), aiming to preserve local variations of gradients in the spatial domain as much as possible, thereby yielding higher-resolution, class-specific attribution maps. This yielded a 3D saliency map for every trial. We chose, among the nine trial-averaging settings, the model that achieved the highest mean validation accuracy as the target model for subsequent Guided Grad-CAM visualization and statistical inference. All visualizations were generated using an independent test set. Note that, to prevent information leakage and selection bias, this test set was used exclusively for visualization and evaluation and was not involved in any model selection or hyperparameter tuning.

Statistical evaluation was performed on correct test trials. Specifically, we organized the Guided Grad-CAM saliency maps for the small-object and large-object conditions into separate four-dimensional arrays (trials × *X* × *Y* × *Z*). We then performed voxel-wise one-sample t-tests against zero. Based on these t-statistics, we then conducted permutation-based cluster-extent inference (5,000 permutations)(Nichols & Holmes, 2002; Winkler et al., 2014). We first utilized the cluster-forming threshold of *P* < 0.05 to identify spatially contiguous, suprathreshold voxel clusters. Then, at the cluster level, we applied an FWE-corrected significance threshold of *P* < 0.05 and retained only clusters surviving this threshold as regions carrying reliable Guided Grad-CAM signals under the small-object and large-object conditions.

In addition, to quantify the key information-bearing regions of the deep model at the anatomical level, we defined 110 regions of interest (ROIs) based on the Harvard–Oxford cortical and subcortical structural atlases. Note that these ROIs were bilaterally divided and screened to exclude non–gray-matter regions such as white matter and ventricles. The mean Guided Grad-CAM value within each ROI mask was computed, resulting in a “trials × ROIs” matrix of average saliency values. Then, one-sample Wilcoxon signed-rank tests were performed on the mean Guided Grad-CAM values for each ROI separately in the small-object and large-object conditions to assess whether they were significantly greater than zero. All ROI-level *P*-values were corrected for multiple comparisons using FDR (*q* < 0.05). If an ROI reached significance in a given condition, this indicates that its mean Guided Grad-CAM value was reliably positive under that condition, reflecting a stable positive attribution of that region to the model’s output for the corresponding class.

## Data availability

All custom Python codes as well as multimodal data are available at https://github.com/masaki-takeda/dld.

## Author Contributions

K.M. and M.T. conceived the project. M.T. collected data. C.L. analyzed data. C.L., K.M. and M.T. wrote the manuscript.

## Acknowledgments

We would like to thank Ms. Maoko Yamanaka for her administrative assistance. This study was supported by KAKENHI from Japan Society for the Promotion of Science (20H00521 and 21K18267 to M.T.), Adaptable and Seamless Technology Transfer Program through Target-driven R&D (A-STEP) from Japan Science and Technology Agency (JST) Grant Number JPMJTR25UE to M.T., a grant from Takeda Science Foundation to M.T., and a grant from Uehara Memorial Foundation to M.T.

