## Supporting Information for "Deep-learning-based fMRI decoding of real-world size for hand-held objects"

*: Correspondence: Masaki Takeda

**This file includes:**

Figure S1 to S2, Table S1 to S2

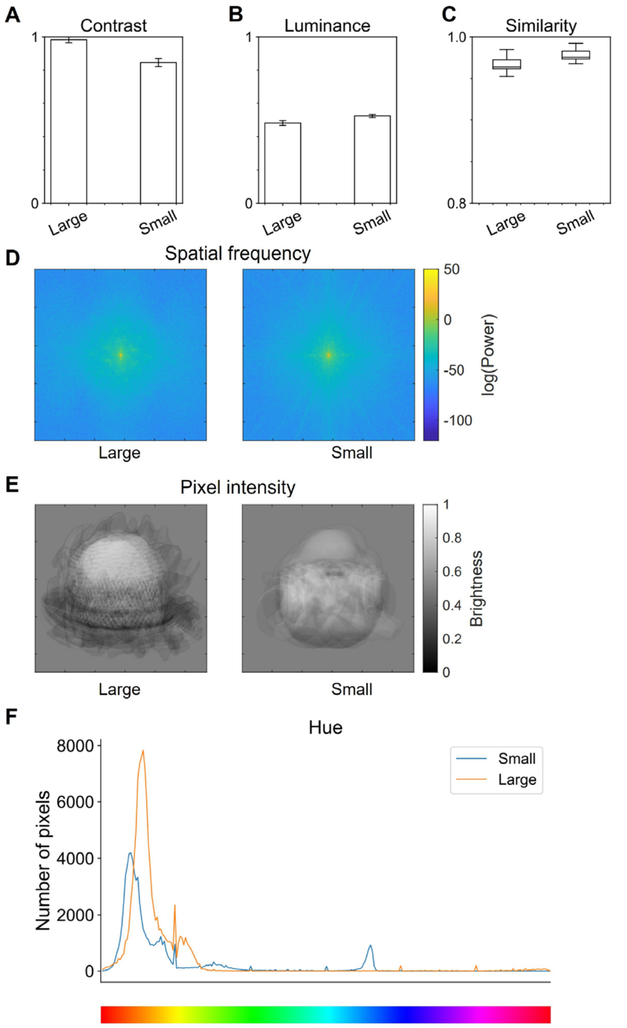


**Figure S1. Low-level visual features of stimulus images.** (A) Michelson contrast of images as the ratio between the difference and the sum of the maximum and minimum pixel intensities. (B) Luminance as the mean pixel intensity of images. (C) Pixel-based pairwise similarity. To estimate the visual similarity among the images in a category, the mean Euclidean distance between normalized grayscale values (scaled to be between 0 and 1) of each stimulus and every other stimulus of the same subcategory was measured. The pairwise similarity values range from 0 (images with inverted intensity difference at each pixel) to 1 (identical images). Means ± S.D. are shown in (A–C). (D) Average spatial frequency across images. (E) Average pixel intensity for each pixel. (F) Average hue histogram across images.

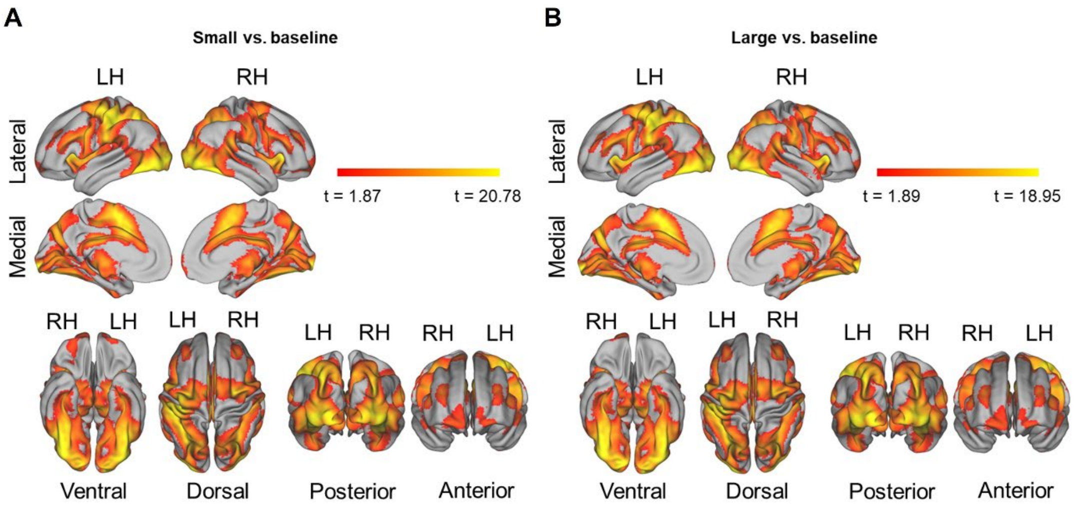


**Figure S2. Spatial representation of small and large objects revealed by univariate analysis.** (A) Spatial representation of small objects. Hot colors denote significant clusters exhibiting stronger responses to small objects. (B) Spatial representation of large objects. Hot colors denote significant clusters exhibiting stronger responses to large objects.

**Table S1. Guided Grad-CAM values for small object trials in anatomically-defined brain regions.**

| Label | Region | Value | FDR *q* | | Label | Region | Value | FDR *q* | |
|---|---|---|---|---|---|---|---|---|---|
| R. Lateral Occipital Cortex, superior division | O | 9.99E-05 | 4.02E-49 | * | R. Parahippocampal Gyrus, anterior division | T | 4.75E-06 | 6.17E-18 | * |
| R. Superior Parietal Lobule | P | 9.83E-05 | 1.27E-48 | * | R. Accumbens | S | 4.72E-06 | 2.20E-20 | * |
| L. Inferior Temporal Gyrus, posterior division | T | 8.83E-05 | 1.44E-37 | * | L. Brain-Stem | S | 4.63E-06 | 2.39E-30 | * |
| R. Insular Cortex | S | 5.46E-05 | 4.57E-38 | * | R. Planum Temporale | T | 4.47E-06 | 9.52E-17 | * |
| R. Putamen | S | 4.61E-05 | 1.39E-35 | * | R. Middle Temporal Gyrus, temporooccipital pa | T | 4.33E-06 | 8.48E-31 | * |
| R. Caudate | S | 4.12E-05 | 2.10E-45 | * | R. Temporal Fusiform Cortex, anterior division | T | 4.11E-06 | 1.16E-40 | * |
| R. Supracalcarine Cortex | O | 2.80E-05 | 1.44E-19 | * | L. Precentral Gyrus | F | 4.03E-06 | 7.66E-37 | * |
| L. Temporal Fusiform Cortex, posterior division | T | 2.64E-05 | 4.66E-42 | * | L. Parahippocampal Gyrus, anterior division | T | 3.95E-06 | 7.89E-13 | * |
| L. Parahippocampal Gyrus, posterior division | T | 2.56E-05 | 2.98E-29 | * | L. Middle Temporal Gyrus, anterior division | T | 3.42E-06 | 6.77E-30 | * |
| L. Parietal Operculum Cortex | P | 2.23E-05 | 9.36E-41 | * | L. Cingulate Gyrus, posterior division | S | 3.31E-06 | 1.76E-20 | * |
| R. Pallidum | S | 2.21E-05 | 2.24E-19 | * | L. Cingulate Gyrus, anterior division | S | 3.28E-06 | 3.45E-15 | * |
| R. Occipital Pole | O | 2.19E-05 | 1.04E-49 | * | L. Accumbens | S | 3.22E-06 | 1.39E-06 | * |
| R. Precuneous Cortex | P | 2.18E-05 | 8.70E-50 | * | L. Pallidum | S | 2.90E-06 | 1.17E-22 | * |
| L. Supracalcarine Cortex | O | 2.04E-05 | 1.85E-28 | * | L. Inferior Temporal Gyrus, anterior division | T | 2.81E-06 | 3.87E-15 | * |
| R. Hippocampus | S | 1.75E-05 | 8.10E-17 | * | R. Cingulate Gyrus, anterior division | S | 2.78E-06 | 6.22E-25 | * |
| R. Cuneal Cortex | O | 1.71E-05 | 2.39E-24 | * | R. Planum Polare | T | 2.50E-06 | 9.51E-24 | * |
| L. Hippocampus | S | 1.69E-05 | 2.65E-24 | * | R. Precentral Gyrus | F | 2.32E-06 | 2.73E-37 | * |
| R. Heschl's Gyrus (includes H1 and H2) | T | 1.64E-05 | 9.26E-32 | * | L. Putamen | S | 2.28E-06 | 8.04E-31 | * |
| R. Temporal Fusiform Cortex, posterior division | T | 1.63E-05 | 8.59E-30 | * | R. Lateral Occipital Cortex, inferior division | O | 2.09E-06 | 3.83E-12 | * |
| L. Occipital Pole | O | 1.54E-05 | 7.35E-28 | * | R. Brain-Stem | S | 2.07E-06 | 9.51E-41 | * |
| R. Parietal Operculum Cortex | P | 1.53E-05 | 7.46E-45 | * | L. Inferior Frontal Gyrus, pars triangularis | F | 2.07E-06 | 6.66E-16 | * |
| R. Angular Gyrus | P | 1.44E-05 | 1.42E-44 | * | R. Frontal Operculum Cortex | F | 2.06E-06 | 2.02E-10 | * |
| R. Supramarginal Gyrus, posterior division | P | 1.30E-05 | 2.65E-41 | * | L. Precuneous Cortex | P | 2.06E-06 | 1.74E-12 | * |
| R. Juxtapositional Lobule Cortex (Supplementary Motor Cortex) | F | 1.24E-05 | 1.63E-45 | * | L. Insular Cortex | S | 1.80E-06 | 2.13E-18 | * |
| L. Supramarginal Gyrus, anterior division | P | 1.13E-05 | 2.92E-21 | * | L. Middle Frontal Gyrus | F | 1.76E-06 | 1.42E-36 | * |
| L. Intracalcarine Cortex | O | 1.10E-05 | 3.79E-38 | * | R. Amygdala | S | 1.73E-06 | 4.39E-09 | * |
| L. Frontal Pole | F | 1.03E-05 | 9.44E-50 | * | L. Frontal Operculum Cortex | F | 1.64E-06 | 0.0004 | * |
| R. Middle Temporal Gyrus, posterior division | T | 1.02E-05 | 5.79E-41 | * | L. Heschl's Gyrus (includes H1 and H2) | T | 1.61E-06 | 0.0012 | * |
| R. Inferior Temporal Gyrus, posterior division | T | 9.88E-06 | 1.70E-39 | * | R. Occipital Fusiform Gyrus | O | 1.61E-06 | 6.40E-15 | * |
| L. Inferior Temporal Gyrus, temporooccipital part | T | 9.44E-06 | 4.09E-29 | * | L. Temporal Fusiform Cortex, anterior division | T | 1.48E-06 | 1.71E-20 | * |
| R. Inferior Temporal Gyrus, temporooccipital part | T | 9.44E-06 | 2.33E-48 | * | L. Planum Polare | T | 1.47E-06 | 1.12E-16 | * |
| R. Superior Temporal Gyrus, posterior division | T | 9.27E-06 | 3.51E-39 | * | L. Frontal Medial Cortex | F | 1.45E-06 | 4.24E-10 | * |
| R. Postcentral Gyrus | P | 8.57E-06 | 3.61E-43 | * | L. Lateral Occipital Cortex, inferior division | O | 1.38E-06 | 1.49E-09 | * |
| R. Temporal Occipital Fusiform Cortex | T | 8.51E-06 | 1.95E-42 | * | R. Subcallosal Cortex | S | 1.14E-06 | 7.04E-19 | * |
| L. Lingual Gyrus | O | 8.49E-06 | 3.28E-33 | * | L. Frontal Orbital Cortex | F | 1.05E-06 | 0.0013 | * |
| L. Temporal Occipital Fusiform Cortex | T | 8.38E-06 | 1.13E-40 | * | R. Frontal Orbital Cortex | F | 9.56E-07 | 1.85E-06 | * |
| L. Planum Temporale | T | 8.30E-06 | 4.52E-25 | * | L. Occipital Fusiform Gyrus | O | 8.82E-07 | 5.09E-11 | * |
| L. Juxtapositional Lobule Cortex (Supplementary Motor Cortex) | F | 8.20E-06 | 2.23E-13 | * | R. Inferior Temporal Gyrus, anterior division | T | 6.10E-07 | 3.74E-23 | * |
| L. Cuneal Cortex | O | 7.86E-06 | 0.6089 | | L. Superior Temporal Gyrus, anterior division | T | 5.68E-07 | 5.53E-12 | * |
| L. Superior Frontal Gyrus | F | 7.50E-06 | 1.47E-46 | * | L. Subcallosal Cortex | S | 5.32E-07 | 0.0539 | |
| R. Lingual Gyrus | O | 7.39E-06 | 8.04E-20 | * | R. Temporal Pole | T | 4.34E-07 | 1.59E-14 | * |
| L. Postcentral Gyrus | P | 7.05E-06 | 1.48E-48 | * | L. Temporal Pole | T | 3.89E-07 | 5.30E-06 | * |
| R. Thalamus | S | 6.94E-06 | 2.63E-21 | * | R. Superior Frontal Gyrus | F | 3.31E-07 | 1.40E-14 | * |
| L. Angular Gyrus | P | 6.90E-06 | 4.01E-45 | * | R. Inferior Frontal Gyrus, pars opercularis | F | 3.26E-07 | 7.42E-14 | * |
| L. Caudate | S | 6.79E-06 | 9.16E-09 | * | L. Amygdala | S | 3.09E-07 | 0.0446 | * |
| L. Supramarginal Gyrus, posterior division | P | 6.33E-06 | 3.58E-49 | * | R. Superior Temporal Gyrus, anterior division | T | 2.16E-07 | 5.42E-05 | * |
| L. Superior Parietal Lobule | P | 6.25E-06 | 5.01E-49 | * | R. Middle Temporal Gyrus, anterior division | T | 1.35E-07 | 0.0002 | * |
| L. Middle Temporal Gyrus, temporooccipital part | T | 6.05E-06 | 5.68E-34 | * | R. Frontal Pole | F | 1.35E-07 | 0.0002 | * |
| R. Cingulate Gyrus, posterior division | S | 5.87E-06 | 1.72E-43 | * | R. Middle Frontal Gyrus | F | 5.25E-08 | 2.72E-05 | * |
| R. Parahippocampal Gyrus, posterior division | T | 5.85E-06 | 1.28E-10 | * | R. Inferior Frontal Gyrus, pars triangularis | F | 2.35E-08 | 7.54E-09 | * |
| R. Intracalcarine Cortex | O | 5.35E-06 | 0.0742 | | R. Frontal Medial Cortex | F | -2.21E-08 | 0.0519 | |
| R. Supramarginal Gyrus, anterior division | P | 5.28E-06 | 5.80E-32 | * | L. Middle Temporal Gyrus, posterior division | T | -3.97E-07 | 0.7080 | |
| L. Central Opercular Cortex | F | 5.26E-06 | 5.04E-37 | * | L. Inferior Frontal Gyrus, pars opercularis | F | -4.02E-07 | 0.4921 | |
| R. Central Opercular Cortex | F | 5.10E-06 | 3.91E-14 | * | L. Superior Temporal Gyrus, posterior division | T | -6.26E-07 | 0.0002 | |
| L. Thalamus | S | 4.89E-06 | 5.19E-18 | * | L. Lateral Occipital Cortex, superior division | O | -1.04E-05 | 5.99E-36 | |

Values are sorted by their magnitudes. An asterisk denotes FDR q < 0.05. O, Occipital; F, Frontal; P, Parietal; T, Temporal; S, Subcortical.

**Table S2. Guided Grad-CAM values for large object trials in anatomically-defined brain regions.**

| Label | Region | Value | FDR *q* | | Label | Region | Value | FDR *q* | |
|---|---|---|---|---|---|---|---|---|---|
| L. Occipital Pole | O | 0.0025 | 5.33E-33 | * | L. Cingulate Gyrus, anterior division | S | 5.85E-07 | 1.18E-09 | * |
| R. Occipital Fusiform Gyrus | O | 1.93E-05 | 5.12E-28 | * | R. Caudate | S | 5.70E-07 | 1.84E-11 | * |
| R. Accumbens | S | 9.32E-06 | 9.58E-12 | * | R. Temporal Pole | T | 5.31E-07 | 5.81E-14 | * |
| R. Thalamus | S | 8.56E-06 | 3.46E-32 | * | L. Pallidum | S | 4.72E-07 | 0.0110 | * |
| R. Pallidum | S | 8.51E-06 | 7.39E-30 | * | R. Cingulate Gyrus, posterior division | S | 4.47E-07 | 2.87E-09 | * |
| R. Lingual Gyrus | O | 8.26E-06 | 3.85E-32 | * | R. Angular Gyrus | P | 4.44E-07 | 9.02E-10 | * |
| R. Brain-Stem | S | 7.33E-06 | 2.16E-24 | * | L. Supracalcarine Cortex | O | 4.43E-07 | 8.51E-08 | * |
| L. Temporal Occipital Fusiform Cortex | T | 5.69E-06 | 3.18E-28 | * | L. Middle Temporal Gyrus, temporooccipital part | T | 4.15E-07 | 1.90E-21 | * |
| R. Subcallosal Cortex | S | 5.36E-06 | 7.61E-11 | * | L. Superior Temporal Gyrus, anterior division | T | 4.00E-07 | 0.4669 | |
| L. Precentral Gyrus | F | 5.06E-06 | 6.46E-30 | * | R. Superior Temporal Gyrus, posterior division | T | 3.75E-07 | 0.0019 | * |
| L. Subcallosal Cortex | S | 4.98E-06 | 1.34E-24 | * | R. Insular Cortex | S | 3.74E-07 | 2.09E-11 | * |
| L. Accumbens | S | 4.52E-06 | 6.02E-18 | * | R. Parahippocampal Gyrus, anterior division | T | 3.71E-07 | 6.82E-08 | * |
| R. Lateral Occipital Cortex, inferior division | O | 4.04E-06 | 5.43E-07 | * | R. Temporal Fusiform Cortex, posterior division | T | 3.47E-07 | 4.49E-15 | * |
| L. Superior Parietal Lobule | P | 3.79E-06 | 1.02E-11 | * | R. Superior Frontal Gyrus | F | 3.45E-07 | 3.11E-13 | * |
| R. Putamen | S | 3.73E-06 | 5.06E-24 | * | R. Heschl's Gyrus (includes H1 and H2) | T | 2.32E-07 | 1.06E-05 | * |
| L. Brain-Stem | S | 3.33E-06 | 2.40E-07 | * | L. Frontal Medial Cortex | F | 2.24E-07 | 7.74E-07 | * |
| R. Cuneal Cortex | O | 3.19E-06 | 0.6779 | | R. Frontal Medial Cortex | F | 2.06E-07 | 1.15E-18 | * |
| L. Thalamus | S | 3.02E-06 | 1.26E-10 | * | R. Middle Temporal Gyrus, posterior division | T | 2.00E-07 | 0.1332 | |
| L. Frontal Orbital Cortex | F | 2.91E-06 | 3.11E-26 | * | R. Middle Temporal Gyrus, anterior division | T | 1.87E-07 | 6.72E-17 | * |
| R. Parahippocampal Gyrus, posterior division | T | 2.67E-06 | 6.38E-27 | * | R. Middle Temporal Gyrus, temporooccipital part | T | 1.86E-07 | 0.0142 | * |
| L. Caudate | S | 2.56E-06 | 1.42E-18 | * | L. Amygdala | S | 1.78E-07 | 0.0001 | * |
| L. Inferior Temporal Gyrus, temporooccipital part | T | 2.49E-06 | 4.61E-29 | * | R. Parietal Operculum Cortex | P | 1.23E-07 | 0.0980 | |
| L. Middle Frontal Gyrus | F | 2.31E-06 | 2.42E-23 | * | R. Central Opercular Cortex | F | 9.98E-08 | 7.46E-05 | * |
| L. Postcentral Gyrus | P | 2.29E-06 | 5.12E-28 | * | R. Middle Frontal Gyrus | F | 9.96E-08 | 5.55E-08 | * |
| L. Inferior Frontal Gyrus, pars triangularis | F | 2.27E-06 | 4.76E-09 | * | L. Middle Temporal Gyrus, anterior division | T | 9.26E-08 | 0.0591 | |
| L. Supramarginal Gyrus, anterior division | P | 2.20E-06 | 4.37E-05 | * | R. Planum Temporale | T | 8.53E-08 | 0.9422 | |
| L. Inferior Frontal Gyrus, pars opercularis | F | 2.19E-06 | 1.64E-15 | * | L. Central Opercular Cortex | F | 7.95E-08 | 0.6986 | |
| L. Middle Temporal Gyrus, posterior division | T | 1.86E-06 | 9.08E-07 | * | R. Juxtapositional Lobule Cortex (Supplementary Motor Cortex) | F | 7.92E-08 | 0.0111 | * |
| R. Occipital Pole | O | 1.81E-06 | 0.4200 | | L. Inferior Temporal Gyrus, anterior division | T | 7.56E-08 | 0.0114 | * |
| L. Precuneous Cortex | P | 1.79E-06 | 1.83E-20 | * | R. Superior Parietal Lobule | P | 6.64E-08 | 2.61E-16 | * |
| R. Precentral Gyrus | F | 1.71E-06 | 4.84E-16 | * | R. Supramarginal Gyrus, posterior division | P | 4.33E-08 | 5.45E-09 | * |
| L. Cuneal Cortex | O | 1.63E-06 | 7.59E-10 | * | R. Superior Temporal Gyrus, anterior division | T | 3.08E-08 | 0.0078 | * |
| R. Postcentral Gyrus | P | 1.62E-06 | 1.19E-18 | * | R. Supracalcarine Cortex | O | 3.04E-08 | 0.5608 | |
| L. Parahippocampal Gyrus, anterior division | T | 1.44E-06 | 0.1612 | | R. Frontal Operculum Cortex | F | 2.69E-08 | 0.0417 | * |
| R. Hippocampus | S | 1.41E-06 | 3.85E-26 | * | R. Frontal Pole | F | 2.44E-08 | 6.07E-07 | * |
| L. Temporal Pole | T | 1.32E-06 | 6.04E-18 | * | R. Supramarginal Gyrus, anterior division | P | 1.31E-08 | 1.47E-07 | * |
| R. Precuneous Cortex | P | 1.29E-06 | 4.79E-22 | * | R. Inferior Frontal Gyrus, pars opercularis | F | 1.31E-08 | 0.0085 | * |
| L. Superior Temporal Gyrus, posterior division | T | 1.28E-06 | 2.22E-05 | * | R. Inferior Temporal Gyrus, posterior division | T | 1.26E-08 | 0.1223 | |
| L. Superior Frontal Gyrus | F | 1.25E-06 | 8.36E-25 | * | R. Temporal Occipital Fusiform Cortex | T | 9.34E-09 | 0.8903 | |
| R. Planum Polare | T | 1.25E-06 | 6.17E-16 | * | L. Planum Temporale | T | 9.00E-09 | 0.0142 | * |
| L. Temporal Fusiform Cortex, posterior division | T | 1.25E-06 | 0.0002 | * | R. Inferior Frontal Gyrus, pars triangularis | F | 7.97E-09 | 2.19E-05 | * |
| L. Supramarginal Gyrus, posterior division | P | 1.18E-06 | 0.1332 | | R. Inferior Temporal Gyrus, anterior division | T | 6.85E-09 | 3.05E-07 | * |
| R. Lateral Occipital Cortex, superior division | O | 1.15E-06 | 2.90E-13 | * | R. Temporal Fusiform Cortex, anterior division | T | 5.55E-09 | 0.3040 | |
| L. Putamen | S | 1.08E-06 | 9.70E-08 | * | L. Parietal Operculum Cortex | P | 4.82E-09 | 0.0018 | * |
| R. Cingulate Gyrus, anterior division | S | 1.02E-06 | 1.85E-15 | * | R. Inferior Temporal Gyrus, temporooccipital part | T | -1.95E-09 | 0.6986 | |
| L. Frontal Pole | F | 9.75E-07 | 7.15E-23 | * | R. Intracalcarine Cortex | O | -7.68E-08 | 0.6986 | |
| L. Cingulate Gyrus, posterior division | S | 9.65E-07 | 1.81E-16 | * | L. Heschl's Gyrus (includes H1 and H2) | T | -3.28E-07 | 6.20E-09 | |
| L. Lateral Occipital Cortex, superior division | O | 9.38E-07 | 5.55E-08 | * | L. Insular Cortex | S | -3.32E-07 | 0.0011 | |
| L. Juxtapositional Lobule Cortex (Supplementary Motor Cortex) | F | 9.19E-07 | 3.72E-16 | * | R. Amygdala | S | -4.49E-07 | 0.0266 | |
| L. Angular Gyrus | P | 8.90E-07 | 0.0371 | * | L. Hippocampus | S | -5.62E-07 | 9.02E-07 | |
| L. Temporal Fusiform Cortex, anterior division | T | 8.23E-07 | 1.38E-07 | * | L. Planum Polare | T | -1.00E-06 | 6.97E-11 | |
| R. Frontal Orbital Cortex | F | 7.74E-07 | 8.30E-17 | * | L. Intracalcarine Cortex | O | -1.65E-06 | 2.96E-06 | |
| L. Inferior Temporal Gyrus, posterior division | T | 7.32E-07 | 3.00E-17 | * | L. Lingual Gyrus | O | -1.29E-05 | 2.01E-12 | |
| L. Frontal Operculum Cortex | F | 7.17E-07 | 1.33E-06 | * | L. Lateral Occipital Cortex, inferior division | O | -0.0002 | 5.33E-33 | |
| L. Parahippocampal Gyrus, posterior division | T | 6.81E-07 | 0.0002 | * | L. Occipital Fusiform Gyrus | O | -0.0002 | 5.33E-33 | |

Values are sorted by their magnitudes. An asterisk denotes FDR q < 0.05. O, Occipital; F, Frontal; P, Parietal; T, Temporal; S, Subcortical.
